# Cavalcade-Mediated Resistance Alters Tomato–Root-Knot Nematode Interactions and Limits Nematode Infection

**DOI:** 10.64898/2026.05.18.726089

**Authors:** Natthidech Beesa, Tarja Hoffmeyer, Arunee Suwanngam, Laura Villegas, Austine Tweneboah, Anongnuch Sasnarukkit, Mohammed Errbii, Buncha Chinnasri, Philipp H. Schiffer

**Affiliations:** Department of Plant Pathology, Faculty of Agriculture, Kasetsart University, Bangkok, 10900, Thailand; Worm∼lab, Institute for Zoology, University of Cologne, Zülpicher Str. 47B, 50674, Cologne, Germany; School of Agriculture, Meiji University, Kawasaki, Kanagawa 214-8571, Japan; Palaeontology & Geobiology, Department of Earth and Environmental Sciences, Ludwig-Maximilians-University Munich, Richard-Wagner-Str. 10, 80333 Munich, Germany; Biodiversity Genomics Center Cologne, University of Cologne, Zülpicher Str. 47B, 50674, Cologne, Germany

**Keywords:** Defense-related genes, Induced plant defense, Nematode management

## Abstract

*Meloidogyne incognita* is a major plant-parasitic nematode responsible for substantial yield losses in tomato worldwide. Current control strategies rely heavily on chemical nematicides, which raise environmental concerns and face increasing regulatory restrictions, underscoring the need for sustainable alternatives. Here, we show that foliar application of an aqueous extract from cavalcade (*Centrosema pascuorum*) enhances tomato resistance against *M. incognita*. Pre-inoculation treatment with cavalcade extract prior to inoculation with root-knot nematodes (RKN) significantly reduced root gall formation, delayed nematode development, and limited second-stage juvenile penetration compared with untreated infected controls, whereas post-inoculation application conferred partial protection. Transcriptomic analyses revealed the activation of multiple defense-related pathways, including salicylic acid– and jasmonic acid–associated signaling and phenylpropanoid metabolism, supported by the upregulation of *PR1* and *PAL*. Additional induction of lipid transfer proteins, leucine-rich repeat receptor-like kinases, resistance proteins, mitochondrial calcium uniporter, and GA2ox5 suggests coordinated activation of pathogen recognition, calcium signaling, and hormone-regulated defense networks. These findings demonstrate that cavalcade extract primes broad-spectrum defense responses in tomato and highlight its potential as an environmentally sustainable strategy for nematode management.

## Introduction

Root-knot nematodes (*Meloidogyne* spp.; RKNs) are globally distributed plant parasites, infecting a broad range of vascular plants in both controlled cultivation settings and open-field agriculture^1^. They are considered the most damaging group of plant-parasitic nematodes because their root damage impairs water and nutrient uptake, causing significant reductions in crop yield and quality across many economically important crops^2^. The most harmful species include *Meloidogyne arenaria, M. incognita, M. javanica*, and *M. hapla*^3^, with recent additions to this list now recognizing *M. enterolobii* as a significant threat^4^ as well. *M. incognita* is one of the most prevalent RKN species, particularly affecting crops such as cotton, cucumber, and tomato^5-7^. Yield losses attributed to *M. incognita* vary significantly, ranging from 12% to 60% in cucumber^8^ and from 22% to 30% in tomato^6^. The presence of this nematode in agricultural fields poses substantial constraints on production systems, including in many developing countries in the global South, underscoring the urgent need for effective management strategies to mitigate its impact.

Although farmers primarily rely on chemical control methods, these practices can cause serious environmental damage and pose health risks to users^9^. Consequently, several studies have sought alternative approaches to reduce dependence on chemical applications. One successful method relies on the activation of plant defense responses against pathogens using biotic and abiotic inducers^10^. These defense responses involve phytohormones such as salicylic acid (SA), jasmonic acid (JA), and ethylene (ET), which serve as signaling molecules in triggering the immune system^11,12^. These phytohormone-mediated pathways are also known to play important roles in plant responses to plant-parasitic nematodes, including RKNs^13,14^. SA signaling is generally associated with resistance to biotrophic and hemibiotrophic pathogens, whereas resistance to necrotrophic pathogens is primarily activated through the combined signaling of JA and ET^15,16^. In the case of SA, its increased levels trigger Systemic Acquired Resistance (SAR), a long-lasting, broad-spectrum defense mechanism that extends to uninfected parts of the plant. This leads to the production of pathogenesis-related (PR) proteins, including *PR1, PR2*, and *PR5*^16^. Hamamouch, et al. ^17^ provided evidence that elevated *PR-1* expression reduced the success rate of infection by both *Heterodera schachtii* and *M. incognita*. In dicotyledonous plants, the expression of *PR-1* is commonly used as a marker for SAR^18^. In addition to *PR-1*, phenylalanine ammonia-lyase (PAL) is extensively studied in plant biology because it links primary metabolism to the secondary phenylpropanoid pathway, playing a key role in plant growth, development, and stress responses ^19^. The upregulation of PAL is particularly important for SA biosynthesis and the activation of defense-related genes^20,21^.

Several chemical compounds have been reported to induce SAR, including acibenzolar-S-methyl (ASM), benzothiadiazole (BTH), probenazole, methyl jasmonate, chitosan, flavonoids, and coumarins^22-24^. For example, flavonoids are secondary metabolites that act as biostimulants, playing a significant role in plant growth by enhancing resistance to a range of biotic and abiotic stresses^25^. Coumarins trigger plant immune responses by promoting the production of reactive oxygen species (ROS), upregulating immunity-related gene expression, and activating signaling pathways such as the salicylic acid–dependent pathway^24^. In general, plants employ ROS as an initial defense mechanism against nematode invasion^26^. In tomato, JA reduces RKN susceptibility through root exudates, with the flavonol kaempferol identified as a key compound suppressing nematode infection^13^.

Cavalcade (*Centrosema pascuorum* cv. Cavalcade), a tropical forage legume of the family Fabaceae widely cultivated as a cover crop, enhances soil fertility through symbiotic nitrogen fixation mediated by *Rhizobium* spp. and improves soil structure^27^. Beyond these agronomic functions, it has been reported to suppress selected plant-parasitic nematodes^28^. Previous studies demonstrated that extracts of *C. pascuorum* inhibit egg hatching and reduce the viability of second-stage juveniles (J2) of *M. graminicola*, as well as suppress mixed developmental stages of *Hirschmanniella mucronata* ^29^,^30^. These nematicidal effects have been attributed to bioactive secondary metabolites, including flavonoids (e.g., quercetin, rutin and kaempferol) and coumarins^29^. Notably, flavonoids and coumarins are also implicated in plant defense signalling and may contribute to the activation of host immune responses. However, molecular evidence supporting a role for *C. pascuorum* in inducing resistance against RKNs remains limited, and it is unclear whether the observed protective effects arise primarily from direct toxicity to the nematode, activation of host defense mechanisms, or a combination of both. To address this gap, the present study evaluated the ability of an aqueous extract from cavalcade leaves to enhance resistance in tomato plants against *M. incognita* by examining both nematode infection parameters and host defense responses. Nematode penetration, gall formation, and developmental progression were quantified, while qRT-PCR and RNA sequencing were employed to assess the expression of key defense-related genes, including PR-1 and PAL, and to comprehensively profile transcriptomic changes associated with induced resistance.

## Results

### Pre-inoculation treatment with cavalcade extract reduces root-knot nematode infection in tomato

To assess whether cavalcade extract improves tomato resistance to *M. incognita*, we examined nematode infection parameters and plant responses under different treatment conditions. Tomato plants were treated with distilled water, extract alone, nematode inoculation (RKN), or extract application either before (ExBRKN) or after (RKNBEx) nematode inoculation.

At three weeks after treatment, gall numbers, nematode developmental progression, and plant growth parameters were evaluated. Gall numbers were significantly reduced in plants receiving the extract prior to nematode inoculation (ExBRKN), whereas post-inoculation application of the extract (RKNBEx) resulted in a slight reduction that was not statistically significant compared with the RKN-inoculated control. This difference may be related to the timing of application, as the extract could be more effective during the early stages of nematode infection and establishment, while treatment after inoculation may have limited impact once nematodes are established within the roots. In ExBRKN plants, gall numbers were reduced by 43.75% (*p* = 0.00; one-way ANOVA) in Trial 1 and 32.58% (*p* = 0.00) in Trial 2 compared with the RKN-inoculated control (**Fig. 1**). No gall formation was detected in either the distilled water or extract-only treatments across both trials. Moreover, no statistically significant effects on plant growth parameters – including root biomass, plant height, and fresh shoot biomass – were observed across the treatments (**Supplementary Table S1**), although a slight increasing trend was noted in the extract-treated plants.

**Figure 1.**
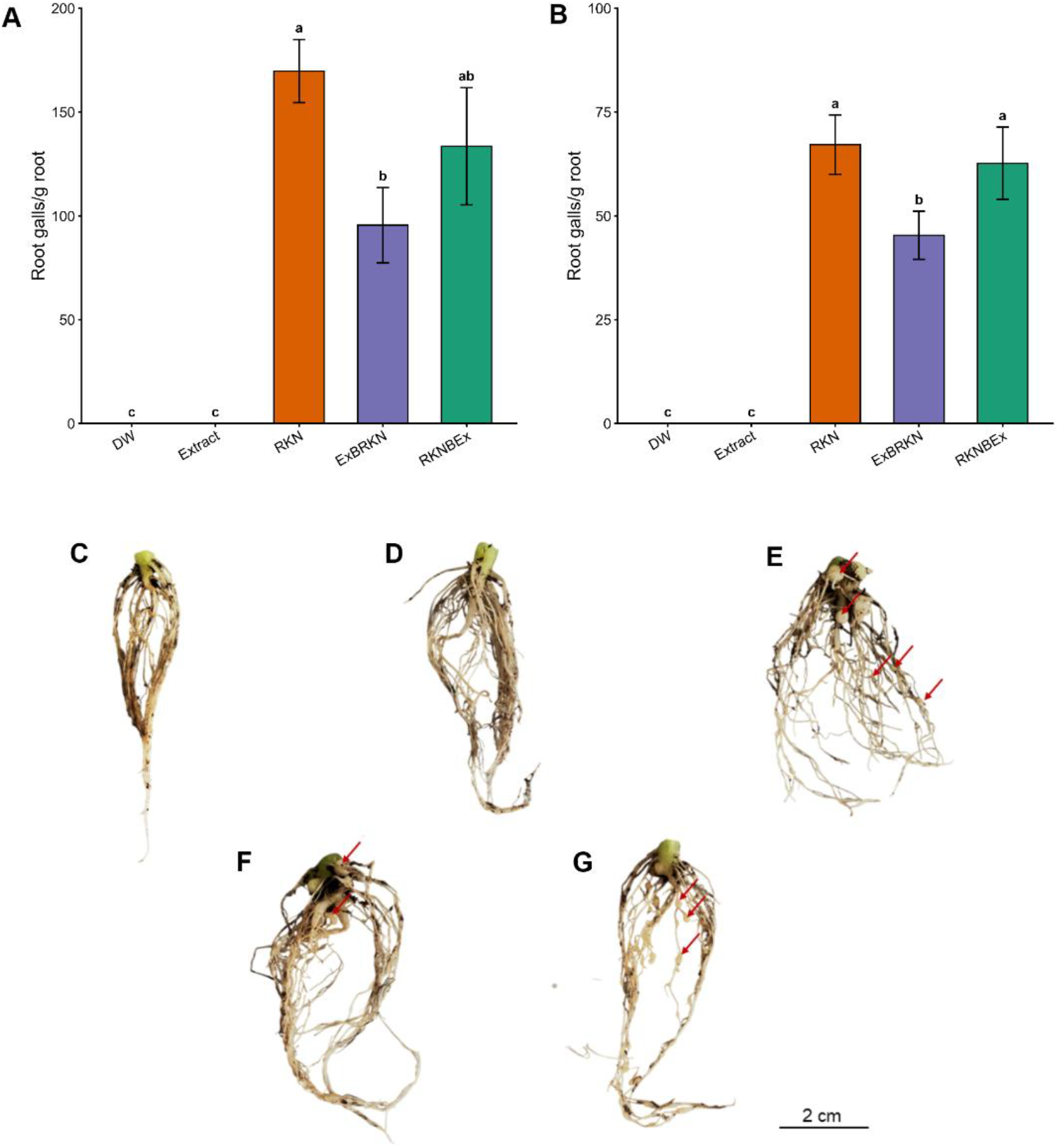
Pre-inoculation application of cavalcade leaf aqueous extract reduces root gall formation in tomato. (A–B) Number of root galls per gram of root tissue in tomato plants subjected to different treatments in two independent experiments (A, Trial 1; B, Trial 2). Data are presented as mean ± S.E.M. (n = 4). Plants inoculated with root-knot nematodes (RKN) showed extensive gall formation, whereas plants pre-treated with cavalcade extract prior to nematode inoculation (ExBRKN) exhibited a significant reduction in gall numbers. Application of the extract after nematode inoculation (RKNBEx) also reduced gall formation, although to a lesser extent than pre-treatment. Distilled water (DW) and extract-only (Extract) controls showed negligible gall formation. Different letters indicate statistically significant differences among treatments within each trial (one-way ANOVA followed by Duncan’s multiple range test (DMRT), p < 0.05). (C–G) Representative root systems of tomato plants under different treatments: DW (C), Extract (D), RKN (E), ExBRKN (F), and RKNBEx (G). Red arrows indicate visible root galls. Scale bar = 2 cm.

In addition to assessing gall formation, whole root systems were stained with acid fuchsin to enumerate nematode developmental stages within the roots. In Trial 1, plants treated with the extract either before (ExBRKN) or after (RKNBEx) nematode inoculation exhibited significantly lower proportions of adult females than the RKN-inoculated control, with reductions of 17.5% and 14.3%, respectively (*p* = 0.00). In contrast, in Trial 2, only the ExBRKN treatment resulted in a significantly lower proportion of adult females than the RKN-inoculated control, with a reduction of 10.9% (*p* = 0.00), whereas the RKNBEx treatment did not differ significantly from the control (**Fig. 2**).

**Figure 2.**
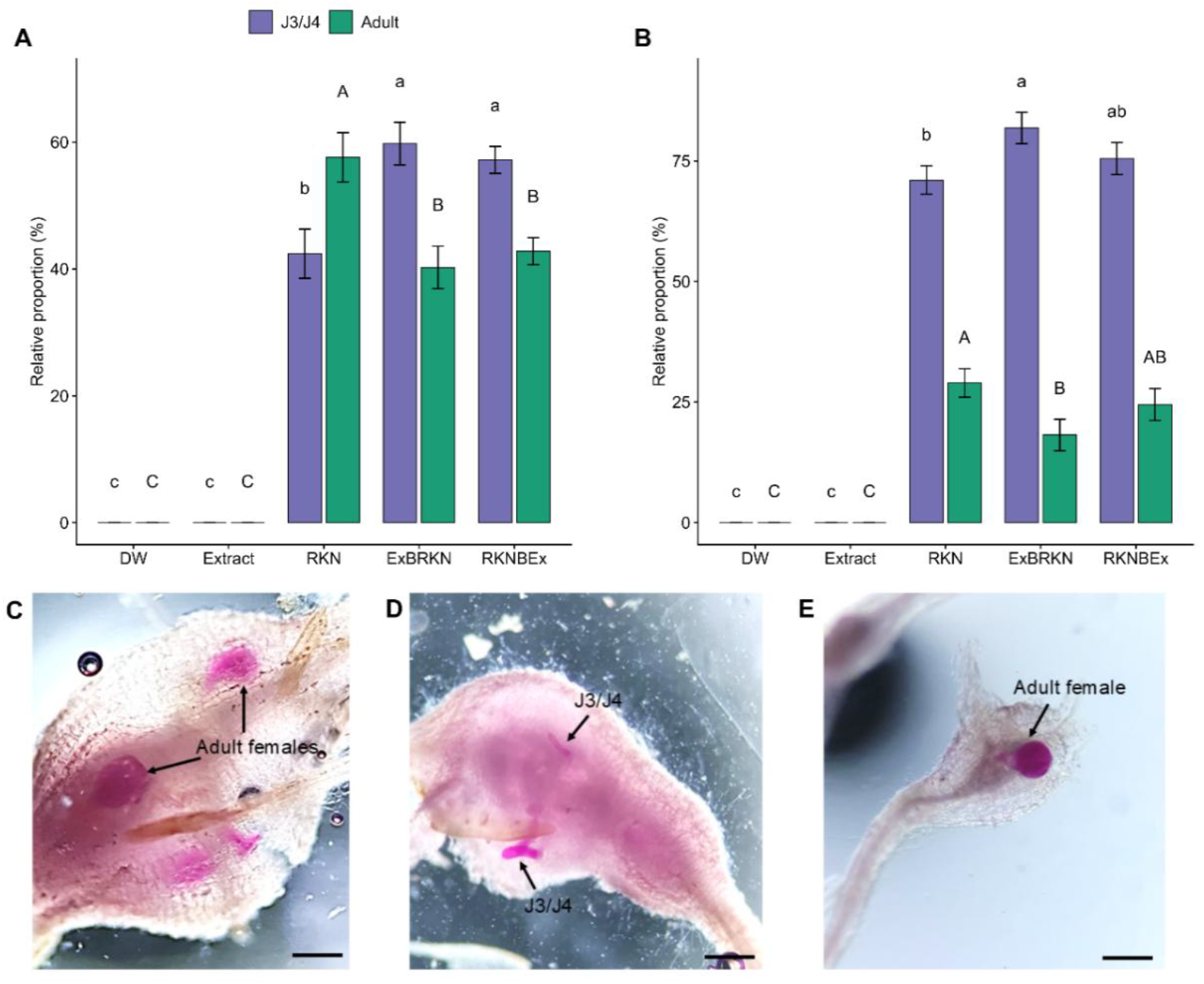
Pre-treatment with cavalcade leaf aqueous extract suppresses the development of *M. incognita* within tomato roots. (A–B) Relative proportions (%) of third- and fourth-stage juveniles (J3/J4) and adult females of *Meloidogyne incognita* recovered from tomato roots under different treatments. Proportions were calculated as the percentage of each developmental stage relative to the total number of nematodes observed per root sample. Tomato plants were treated with cavalcade leaf aqueous extract either before nematode inoculation (ExBRKN) or after inoculation (RKNBEx). Control treatments included nematodes alone (RKN), extract alone, and distilled water (DW). (A) Trial 1; (B) Trial 2. In Trial 1, the mean total numbers of nematodes per gram of root were 213.3 (RKN), 152.6 (ExBRKN), and 170.4 (RKNBEx). In Trial 2, the corresponding values were 90.5, 53.4, and 81.5, respectively. Data represent two independent experiments. Different lowercase letters indicate significant differences among treatments for J3/J4, whereas different uppercase letters indicate significant differences among treatments for adult females, according to Duncan’s multiple range test (*p* ≤ 0.05). (C–E) Representative acid fuchsin–stained tomato roots showing developmental stages of *M. incognita* under different treatments: (C) RKN, (D) ExBRKN, and (E) RKNBEx. Arrows indicate stained nematodes. Scale bars = 500 µm.

To investigate whether foliar application of the extract influences early nematode infection, particularly at the infective J2, a separate experiment was conducted to quantify J2 penetration in tomato roots. At 1 day after treatment, ExBRKN plants exhibited the lowest J2 counts (mean ± SE; 85.0 ± 15.5), whereas RKNBEx plants (341.95 ± 30.22) harbored more J2s than the RKN-inoculated control (165.0 ± 21.9) (*p* = 0.00). By 2 and 3 days after treatment, both ExBRKN (154.8 ± 19.3 and 115.2 ± 23.9) and RKNBEx (293.4 ± 46.3 and 210.6 ± 20.5) displayed significant reductions in J2 infection relative to the RKN-inoculated control (411.9 ± 57.3 and 315.5 ± 29.0) (*p* = 0.00) (**Fig. 3**).

**Figure 3.**
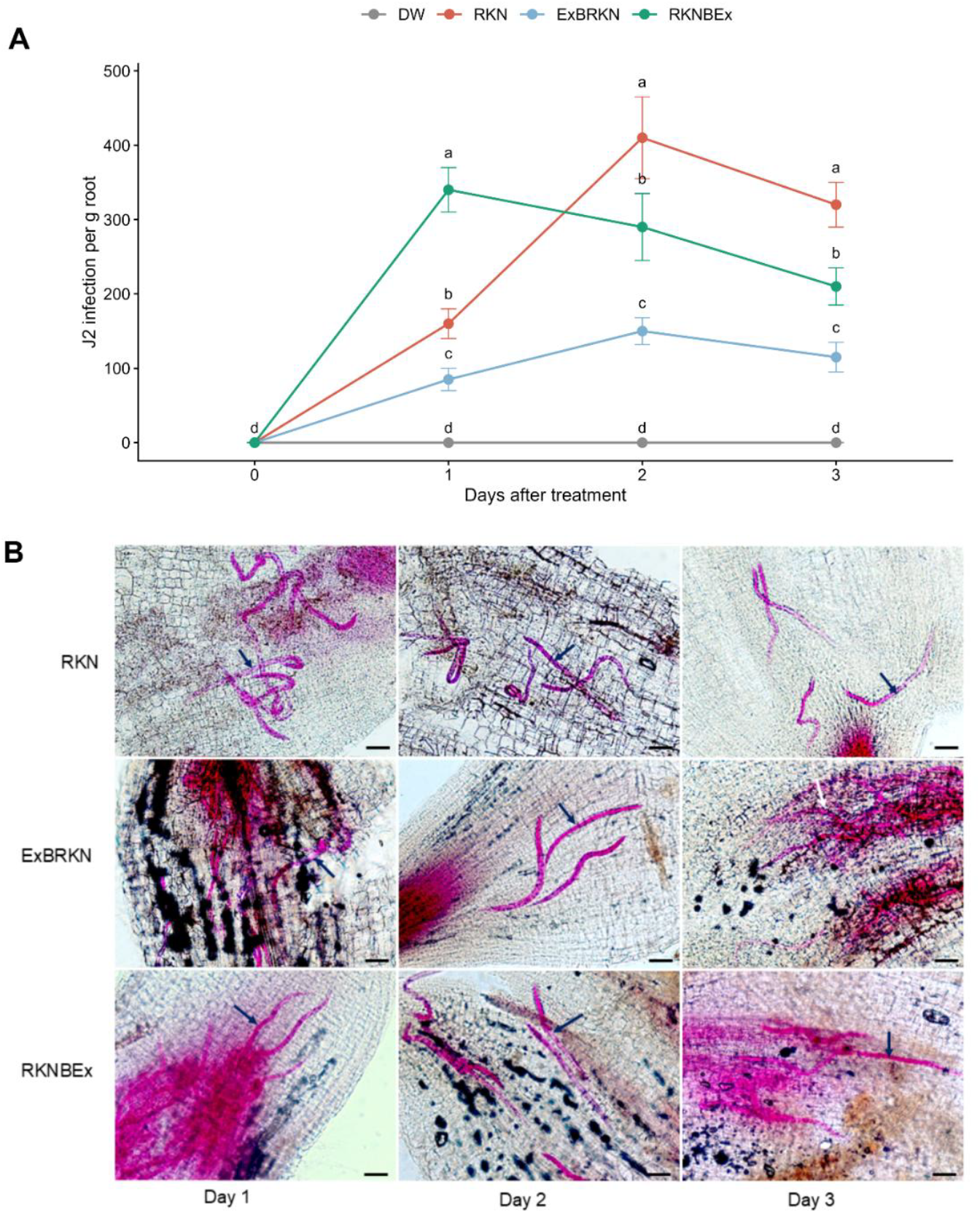
Pre-treatment with cavalcade leaf aqueous extract reduced early infection of *M. incognita* second-stage juveniles (J2) in tomato roots. (A) Number of J2s invading tomato roots at 0–3 days after treatment in plants receiving the extract prior to nematode inoculation (ExBRKN) or after inoculation (RKNBEx). Control treatments included distilled water (DW) and nematodes alone (RKN). Mean differences were assessed using Duncan’s Multiple Range Test at the 0.05 significance level; points sharing the same lowercase letter within each sampling day are not significantly different. (B) Representative acid fuchsin–stained tomato roots showing J2 infection under different treatments across three days. Rows correspond to treatments (RKN, ExBRKN, and RKNBEx), and columns represent days after treatment (Day 1–Day 3). Arrows indicate stained nematodes (RKN). Scale bars = 50 μm.

### Pre-treatment with cavalcade induces defense-related gene expression as revealed by qRT-PCR

To determine if the observed differences in nematode infection were associated with changes in plant defense responses, we analysed the expression of selected defense-related genes in treated and untreated tomato plants via qRT-PCR. We first focused on well-characterized marker genes, *PAL* and *PR1*, which are commonly used indicators of SA–mediated defense pathways, to assess whether their induction patterns corresponded with the differences observed in J2 penetration and early resistance responses. The ExBRKN treatment was used as the representative induced-resistance condition for these targeted analyses.

Foliar application of the extract appears to prime defense responses in treated plants, as evidenced by the significantly elevated *PR1* transcript levels in plants receiving the extract prior to nematode inoculation (ExBRKN), particularly at days 1 and 2 (4.83- and 8.38-fold, respectively), relative to both the RKN-only and extract-only controls (1.45- and 3.54 -fold, respectively). In comparison to these controls, the distilled-water inoculated negative control showed an increase in PR1 expression. Although this expression was not as high as the one in the ExBRKN treated plant, they were not significantly different according to Duncan’s multiple range test. In addition, no significant differences in *PR1* expression were detected among treatments at day 3 (**Fig. 4A**).

**Figure 4.**
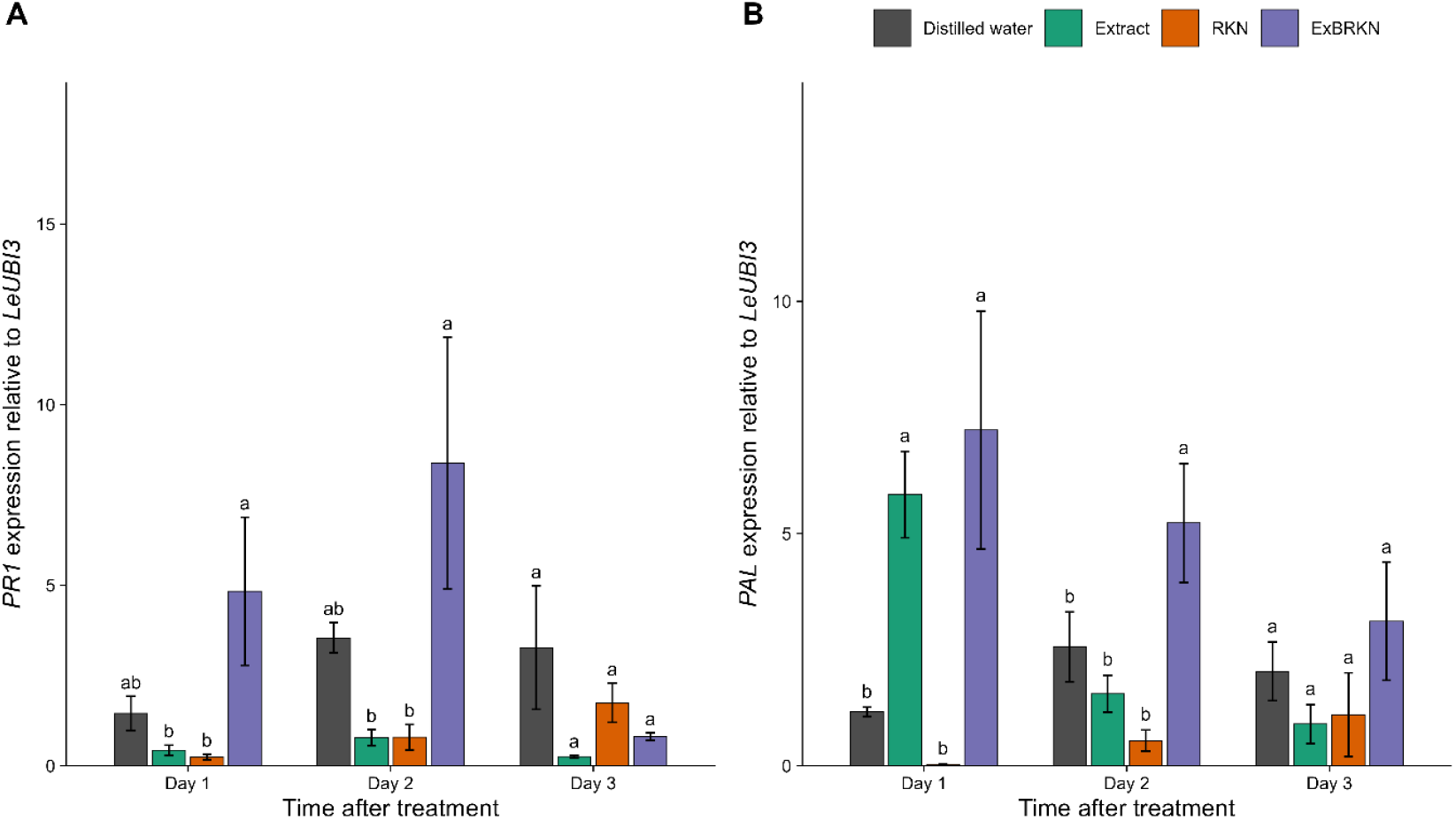
Expression of defense-related genes in tomato roots following extract and nematode treatments, as determined by qRT-PCR. (A) *PR1* and (B) *PAL* transcript levels relative to *LeUBI3* at 1, 2 and 3 days after treatment with distilled water (DW), extract alone, root-knot nematode (RKN), or extract plus RKN (ExBRKN). Bars represent mean ± S.E.M. (n = 3 biological replicates). Different lowercase letters indicate significant differences among treatments within the same time point, as determined by Duncan’s multiple range test (*p* = 0.05).

A similar pattern was observed for *PAL* gene expression, in which the ExBRKN treatment exhibited the highest transcript levels at Days 1 and 2 (7.23- and 5.79 -fold, respectively); however, these increases were not statistically different from the extract-only treatment at Day 1 (5.84-fold). By Day 3, the response had leveled off, and all treatments showed comparable activity in the roots (**Fig. 4B**). Our findings suggest that foliar application of the extract enhances *PAL* gene expression, while its application in combination with subsequent RKN infection enhances *PR1* gene expression. These responses during the early stages of RKN infection, may contribute to the reduction of nematode penetration and subsequent development.

### Pre-treatment with cavalcade induces defense-related gene expression as revealed by transcriptomic analyses

To investigate the physiological response of tomato roots to root-knot nematodes (RKN) in the presence and absence of cavalcade leaf aqueous extract, we performed mRNA sequencing on roots subjected to four treatments: extract applied before RKN inoculation (ExBRKN), extract only (extract), RKN only, and distilled water (DW) as control (n = 3 each).

Of the 41,464 genes annotated in the tomato genome (version SLM_r2.1; NCBI accession: GCF_036512215.1), 15,685 were expressed, and 956 genes showed significant differential expression across treatments (Benjamini–Hochberg adjusted *p* < 0.05; **Table 1**). RKN induced minimal changes in gene expression in the roots compared to the control, with only two genes downregulated, including a transcription factor (LOC101248512) as well as a protein of unknown function (LOC101257056). In contrast, the highest numbers of differentially expressed genes (DEGs) were observed in pairwise comparisons involving ExBRKN (360 genes) or the plant extract (278 genes) versus the control, as well as in RKN versus ExBRKN (311 genes) (**Table 1**).

**Table 1.** Differentially expressed genes between treatments giving the number of expressed genes that were not significantly different, upregulated, or downregulated in each pairwise comparison.

| Comparison | Not Significant | Up regulated | Down regulated |
| --- | --- | --- | --- |
| ExBRKN vs DW | 15325 | 172 | 188 |
| Extract vs DW | 15407 | 226 | 52 |
| RKN vs DW | 15683 | 0 | 2 |
| Extract vs ExBRKN | 15681 | 4 | 0 |
| RKN vs Extract | 15684 | 0 | 1 |
| RKN vs ExBRKN | 15374 | 210 | 101 |

Gene Ontology (GO) analyses of DEGs between ExBRKN and the controls, the plant extract and the controls, and between ExBRKN and RKN shows that the cavalcade leaf extract induces a classic defense-priming program in tomato roots (FDR-adjusted *p* < 0.05; **Supplementary Tables S2-S4**). Enriched GO terms included ethylene-activated signaling pathway (GO:0009873), jasmonic acid biosynthetic process (GO:0009695), regulation of salicylic acid biosynthetic process (GO:0080142), calcium ion transmembrane transport (GO:0070588), and cell wall biogenesis (GO:0042546).

When the extract is applied before nematode inoculation (ExBRKN vs DW) the most significantly enriched functions include zinc-ion transport, ethylene-activated signaling, and DNA-binding transcription-factor activity. In contrast, genes associated with xyloglucan metabolism, cell-wall biogenesis, and heat-shock/chaperone activity are downregulated, suggesting a temporary shift away from growth-related wall remodeling toward defense signaling and barrier formation. The extract alone (Extract vs DW) showed enrichment of genes associated with jasmonic-acid biosynthesis, lipid-metabolism and transcriptional regulation, while genes related to the general heat-shock response are suppressed, indicating a defense-related hormonal response even in the absence of the pathogen.

In the comparison between extract-pre-treated and nematode-only roots (ExBRKN vs RKN), enriched functions include calcium-signaling components (e.g., P-type Ca^2+^ transporters, calmodulin-binding proteins, Ca^2+^ transmembrane transport) together with ATP -dependent chaperones, reflecting activation of early defense signaling. Cell-wall remodeling genes (xyloglucan transferases, cell-wall biogenesis) and actin-nucleation factors are also up-regulated, which may contribute to reinforcement of the physical barrier and cytoskeletal reorganization that may hinder feeding-site formation. Importantly, both salicylic-acid and jasmonic-acid signaling pathways are enriched, indicating a dual-hormone primed state, while apoplastic proteins are induced, providing an enhanced extracellular defense layer.

A multidimensional scaling of the expression data revealed four discrete clusters (**Fig. 5A**). Dimension (Dim) 1, which accounts for 25 % of the total variance, separates the extract-treated samples (Extract and ExBRKN) from the water control, indicating that the leaf aqueous extract is the principal driver of global transcriptional change. Dim 2 (18 % of variance) reflects the modest within-treatment variability.

**Figure 5.**
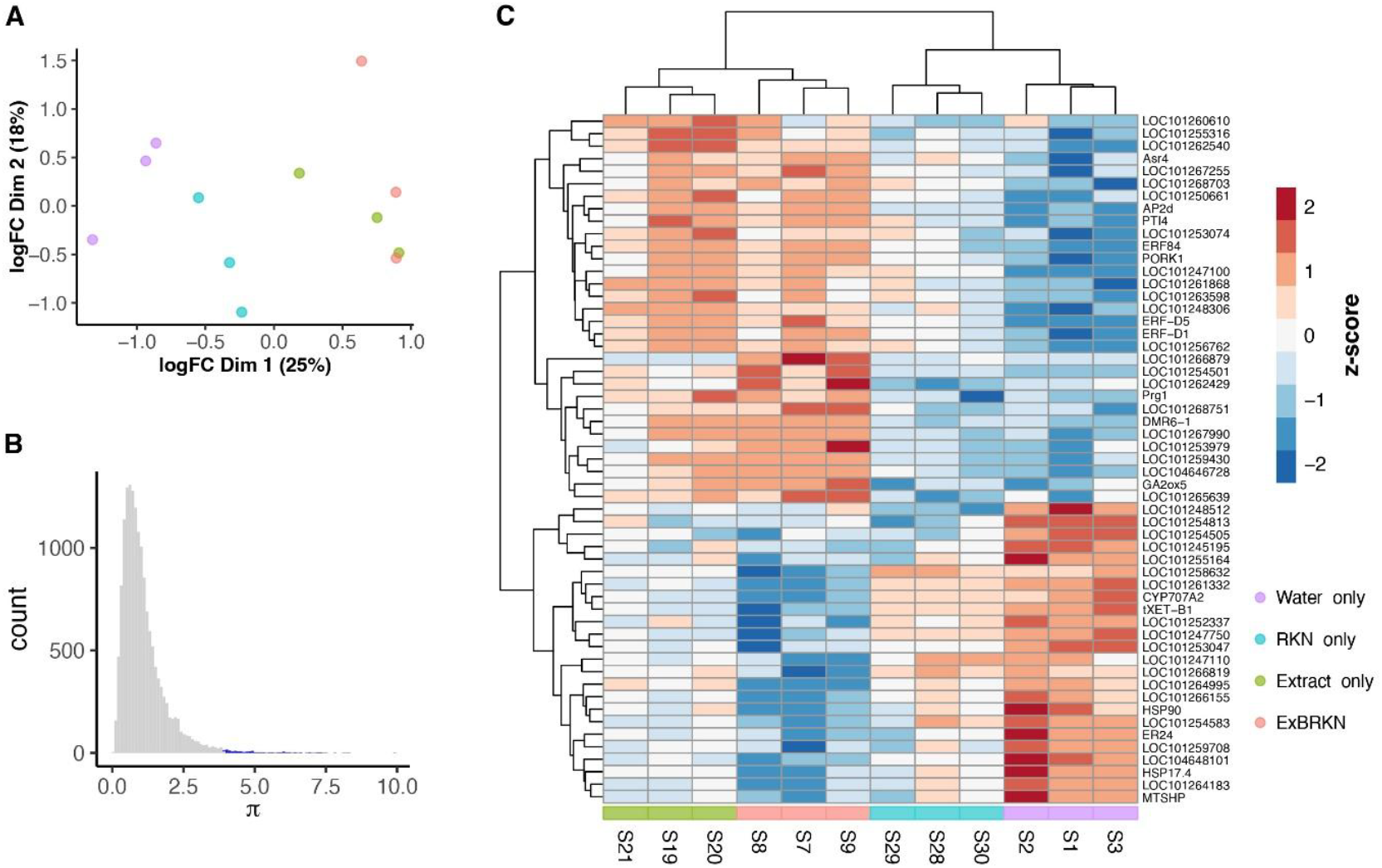
Differential gene expression analysis. (A) Multidimensional scaling (MDS) plot showing separation of treatment groups along Dim1 explaining 25% of the variation. (B) Distribution of plasticity indices (π) quantifying expression bias across treatments with the blue bars indicating the top 1% most biased genes selected for functional analysis. (C) Heatmap of 55 genes with pronounced expression bias among treatments (ExBRKN, extract only, RKN only, and control). Expression values for each gene (rows) are Z-score normalized across samples (columns).

To identify genes contributing to treatment-specific responses, we computed the plasticity index (π), which summarizes expression bias of each gene across multiple contrasts (Schrader et al. 2017). High π values indicate treatment-biased gene expression, whereas low values indicate uniform expression across treatments. We then selected the top 1% most biased genes (**Fig. 5B**) as outliers for further functional analyses, yielding 52 genes with pronounced expression bias. A subset of these genes was upregulated in the roots of plants exposed to the cavalcade extract and ExBRKN treatments, while another subset was downregulated in these treatments compared with roots exposed to RKN only and the controls (**Fig. 5C**).

To infer the putative functions associated with the 52 genes with the highest π, we annotated them using eggNOG-mapper (version 2.1.12)^31^. The resulting annotations (**Supplementary Table S5**) revealed that several genes upregulated in tomato plants are potentially involved in stress responses, secondary metabolism, and cell wall modification, processes likely involved in the plant’s defense response to root knot nematodes. For example, among the most strongly upregulated genes in the ExBRKN treatment, we detected one encoding a putative pathogenesis-related protein, a glucan endo-1,3-beta-glucosidase potentially involved in cell wall degradation, and a cytochrome P450 monooxygenase implicated in secondary metabolite biosynthesis. In contrast, in the RKN-only treatment, the expression levels of these genes were reduced. This contrasting expression pattern suggests that nematode infection alone and in combination with plant extract may differentially influence host gene expression.

## Discussion

*Meloidogyne incognita* is one of the most destructive plant-parasitic nematodes, causing substantial yield losses by disrupting root structure and function in a wide range of host plants. Its severe impact on tomato cultivation and other crops worldwide highlights the critical need for effective and sustainable management strategies^32,33^. Previous studies have shown that cavalcade legume can suppress plant-parasitic nematodes both directly, by killing juveniles and inhibiting hatching, and indirectly, by influencing soil microbial community composition in a way that favors antagonistic microorganisms and contributes to plant - parasitic nematode suppression^29,30^. Cavalcade leaves contain several bioactive compounds, including quercetin, rutin, kaempferol, and coumarins, some of which have been reported to stimulate plant resistance against nematode infection^34,35^. In this study, we evaluated the efficacy of foliar application of cavalcade leaf aqueous extract and subsequently analysed its effects on both the nematode (parasite) and the host plant.

Our data suggest the activation of induced systemic resistance in tomato following treatment with cavalcade extract, which may contribute to the observed suppression of nematode infection. Consistent with this interpretation, the significant reduction in root gall formation and the delayed development of *M. incognita* were accompanied by the marked upregulation of the defense-related genes *PR1* and *PAL* in pre-treated plants under nematode stress. These genes are well-established molecular markers of plant immune activation^36-38^. The increased expression of *PR1* is associated with the activation of the salicylic acid–dependent defense pathway, which plays a pivotal role in systemic acquired resistance against biotrophic pathogens and plant-parasitic nematodes^18^. Consistently, several previous studies have reported that *PR1* is associated with enhanced plant resistance against plant-parasitic nematodes^17,39^. For example, Hamamouch, et al. ^17^ showed in *Arabidopsis thaliana* that elevated *PR-1* expression reduced the infection success of both *H. schachtii* and *M. incognita*, possibly by inducing the production of salicylic acid. Similarly, Charehgani, et al. ^40^ reported that exogenous application of defense-related signaling compounds, including SA, was accompanied by increased expression of *PR1* and other defense-associated genes in tomato, together with reduced nematode infection parameters. In addition to *PR1*, upregulation of *PAL* is commonly associated with activation of the phenylpropanoid pathway, which contributes to the biosynthesis of defense-related secondary metabolites and structural barriers^41-43^. The activation of these pathways likely underlies the reduced gall formation, delayed nematode development, and decreased J2 penetration observed in the present study, confirming that cavalcade extract enhances tomato resistance likely through the coordinated regulation of SA signaling and PAL-mediated phenylpropanoid metabolism.

To further elucidate the molecular mechanisms underlying *PR1*- and *PAL*-mediated resistance, transcriptomic analysis using mRNA sequencing was performed to characterize global defense-related gene expression in tomato following cavalcade extract treatment. The results revealed that, under nematode infection stress, foliar application of the extract significantly induced the expression of multiple resistance-related genes. For example, lipid transfer proteins (LTPs; Gene ID: LOC101266879) were significantly upregulated following cavalcade extract treatment. LTPs are critical for cuticular lipid deposition and contribute to plant defense^44^. In tobacco, elevated LTP expression was associated with increased cuticular wax accumulation^45^, and in *A. thaliana*, LTPg1 has been shown to participate in cuticular lipid accumulation^46,47^. By reinforcing the cuticular barrier, LTPs likely restrict pathogen and nematode invasion. Similarly, a gene encoding a leucine-rich repeat receptor-like protein kinase (LRR-RLK; Gene ID: LOC101262429) was significantly upregulated in tomato roots treated with the extract plus *M. incognita* (ExBRKN) compared with the distilled water control. The upregulation of this gene suggests that cavalcade extract may enhance tomato resistance against *M. incognita* through the activation of pathogen recognition and immune signal transduction. LRR-RLKs act as pattern recognition receptors, detecting pathogen-associated molecular patterns and activating downstream defense signaling^48^. Activation of LRR-mediated signaling triggers early immune responses, including calcium influx, reactive oxygen species production, and the expression of defense genes^49-51^. In addition, resistance proteins (Gene ID: LOC101268751) and genes involved in calcium signaling (Gene ID: LOC101253979) and hormone metabolism (Gene ID: GA2ox5) were also strongly induced. Together, the upregulation of these genes is consistent with the activation of plant defense mechanisms^52-55^, promoting resistance to the RKN *M. incognita*.

Our results suggest that the extract may act as a defense-priming agent, potentially influencing transcriptional regulation and hormone-related defense pathways prior to nematode infection. Upon nematode invasion, this primed state may facilitate a more rapid activation of defense responses, including those associated with SA and JA, together with possible structural reinforcement of root tissues, thereby contributing to the restriction of nematode infection and establishment. These findings are consistent with the report by Martínez-Medina, et al. ^56^, who showed that treatment of tomato plants with the antagonistic fungus *Trichoderma* functions as a defense-priming strategy. In that study, priming of SA-dependent defenses restricted early nematode invasion, whereas activation of JA-dependent defenses reduced feeding-site development and nematode reproduction at later stages of infection^56^. Similarly, Martinuz, et al. ^14^ demonstrated that exogenous application of SA and methyl jasmonate (MeJA) in *A. thaliana* synergistically activated SA- and JA-mediated defense signaling pathways, thereby enhancing resistance against *M. incognita*. Plants treated with the extract alone exhibited activation of JA-related responses, as indicated by the upregulation of genes associated with JA biosynthesis, lipid metabolism, and transcriptional regulation. This response may reflect the perception of bioactive compounds in the extract by plant receptors, which can trigger defense signaling even in the absence of nematode infection^57^. The extract used in this study contains flavonoids, which may contribute to these responses ^28^. Akhtar, et al. ^58^ reported that flavonoid biosynthesis is closely linked to hormonal signaling involved in plant stress responses. In general, abscisic acid (ABA) and JA function as early-responsive hormones under osmotic and ionic stress, whereas auxin and ethylene regulate the balance between plant growth and defense^58^. In contrast, the expression of these defense-related genes was reduced in plants inoculated with nematodes alone. This suppression may be associated with the secretion of effector proteins by *M. incognita*, such as Minc03329, MiEFF1, and MiEFF18, which have been reported to interfere with host immune responses and promote pathogenicity^59-61^.

In summary, foliar application of cavalcade extract appears to strengthen tomato defenses against *M. incognita* through the activation of multiple protective pathways. This is supported by the upregulation of defense-related genes involved in salicylic acid–associated signalling (*PR1*), phenylpropanoid metabolism (*PAL*), and receptor-like kinases linked to pathogen perception. Treated plants showed reduced nematode infection parameters, including fewer galls and egg masses, indicating enhanced resistance. These coordinated responses highlight the potential of cavalcade extract as a safe and sustainable approach for nematode management. Future studies should focus on developing application strategies and exploring its effectiveness in different crops.

## Materials and Methods

### Nematode inoculum preparations

*Meloidogyne incognita* used in the present study was obtained from Dr. Jan Henrik Schmidt at the Institute for Epidemiology and Pathogen Diagnostics, Germany. To verify the nematode species, DNA was extracted from second-stage juveniles using a modified HotSHOT DNA extraction method^62^. The experiment was performed with five replications for validation. DNA was amplified using the Sequence Characterized Amplified Region (SCAR) marker Inc14F/Inc14R^63^, and the resulting 520 bp amplicon confirmed the nematode as *M. incognita* (**Supplementary Figure S1**).

To establish and maintain the nematode inoculum, J2s were introduced into 2-week-old tomato roots grown in 8-cm-diameter plastic pots containing 300 g of autoclaved peat moss. The plants were kept near a laboratory window under a controlled room temperature of 21 °C and were watered three times per week. At two months of age, the infected tomato plants were uprooted and thoroughly rinsed with tap water until clean. Nematode egg masses were picked from the roots under a stereo microscope (ZEISS Stemi 508), placed in a 50 ml Falcon tube containing 30 ml of 0.6% sodium hypochlorite (NaOCl), and shaken by hand for 2 minutes. The nematode egg suspension was then be poured through a 25-µm mesh sieve, rinsed with distilled water until thoroughly clean, and placed on tissue paper lined on a tray. Distilled water was added until the eggs were fully submerged, and the suspension was incubated at room temperature for 7 days. Fresh J2s hatched from the eggs were used in the experiments.

### Extract preparation

Ten grams of ground material (1-month-old cavalcade leaves), prepared according to Beesa, et al. ^28^, were added to 100 ml of distilled water in a 250 ml conical flask and shaken on an incubator shaker (BioSan Shaker-Incubator ES-20) at 150 rpm for 24 hours at 25°C. The mixture was then filtered through muslin cloth, centrifuged at 3,000 × *g* for 10 minutes, and the supernatant was further filtered through 0.45 µm Whatman cellulose acetate membrane filters (Global Life Sciences Solutions Operations, UK). The resulting filtrate (cavalcade extract) was used in subsequent experiments.

### Pot experiments

Gall formation and nematode development: Two-week-old tomato seedlings (*Solanum lycopersicum* ‘Hoffman’s Rentita’) were transplanted into 8-cm-diameter plastic pots containing 300 g of autoclaved peat moss, with one plant per pot. One week later, the following treatments and controls were prepared: 1) plants mock-inoculated with distilled water (DW), 2) plants treated with the extract only (Extract), 3) plants inoculated with RKN at J2 stage only (RKN), 4) plants treated with the extract 24 hours prior to nematode inoculation (ExBRKN), and 5) plants treated with the extract 24 hours after nematode inoculation (RKNBEx). In this study, the extract at 100% concentration was applied to each plant via foliar spraying at a volume of 0.3 ml. Plants were inoculated with a nematode suspension containing 500 J2s in 1 ml per pot. The experiment was arranged in a randomized complete block design (RCBD) with four replications and repeated once. Plants were watered three times per week. Three weeks after treatment, the tomato plants were uprooted and rinsed with tap water until all soil debris was removed. Data were recorded, including plant height (cm), root weight (g), and the number of root galls. Additionally, roots were stained with acid fuchsin to assess the number of developing nematode stages in each treatment, compared with healthy plants and nematode-inoculated controls.

J2 infection: A separate experiment was conducted to investigate the penetration ability of nematode J2s in plants after treatment with the extract. The treatments and experimental conditions followed those of the previous experiment, but plants were collected at 1, 2, and 3 days post-treatment. The experiment was arranged in a randomized complete block design (RCBD) with three replications and repeated once. The collected plants were rinsed with tap water and were stained with acid fuchsin. The number of J2s within the roots were then counted under a stereomicroscope.

### Analysis of defense related genes expression

Tomato plants were subjected to extract and nematode treatments under the same conditions as described in the J2 infection experiment. Root samples were collected at 1, 2, and 3 days after treatment for RNA extraction. For each sample, 0.05 g of root tissue was ground in liquid nitrogen using a mortar and pestle. The resulting powder was transferred to an RNase-free tube containing 450 µL of RLT buffer. Total RNA was then extracted using the RNeasy Mini Kit (Qiagen) according to the manufacturer’s instructions. RNA quality and quantity were determined using a NanoDrop™ 2000c spectrophotometer (Thermo Fisher Scientific).

For each treatment, three biological replicates (individual plants) were analysed, and each biological replicate included three technical replicates. Quantitative RT-PCR (qRT-PCR) was performed using the Luna® Universal One-Step RT-qPCR Kit (Biolab, UK) to quantify the expression of the target genes (*PR1* and *PAL*), with *LeUBI3* used as the internal reference gene, according to the manufacturer’s instructions. Amplification was carried out on a 7500 Fast Real-Time PCR System (Applied Biosystems). Relative expression levels were calculated using 7500 Software v2.3 (Applied Biosystems) based on the ΔΔCt method, in which the Ct values of the target genes were normalized to those of the housekeeping gene LeUBI3^36^. Primer sequences, amplicon sizes and GenBank accession numbers for all analysed genes are listed in **Table 2**.

**Table 2.**
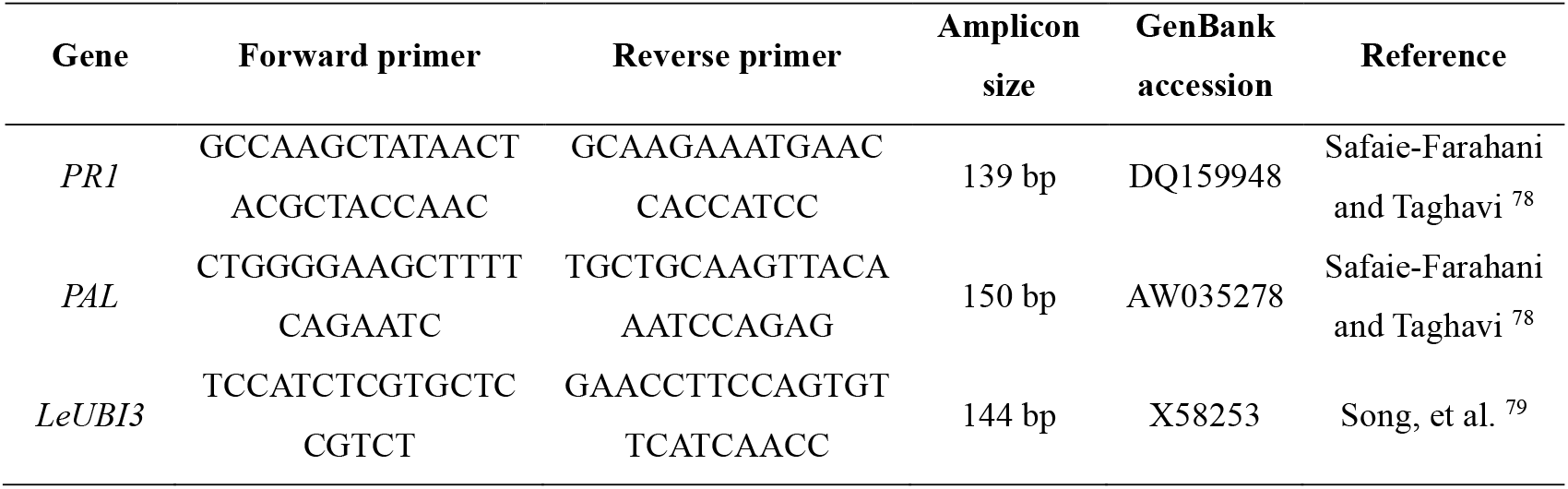
List of primer sequences used for RT–qPCR.

### mRNA sequencing analysis

RNA obtained from all treatments at day 1 after treatment, serving as representative samples, was submitted to Novogene CO., LTD. (Munich, Germany) for mRNA sequencing. RNA sequencing was performed on an Illumina NovaSeq X Plus platform using paired-end 150 bp reads (PE150). Each sample was prepared as an independent mRNA library using poly(A) tail enrichment. A total of approximately 6.4–9.8 million raw reads were generated per sample. The sequencing quality was high, with Q20 > 98% and Q30 > 96%. Transcriptome and gene expression analyses were performed as described previously^64^. Briefly, the raw reads quality was assessed using FastQC [https://www.bioinformatics.babraham.ac.uk/projects/fastqc/] and ribosomal RNA (rRNA) sequences were removed using SortMeRNA ^65^ and the Silva databases (version 138.2) ^66^. Short, low-quality reads and adapter contamination were filtered out using Trimmomatic (version 0.38) ^67^ with the following options: ILLUMINACLIP:TruSeq3-PE.fa:2:30:10:2:keepBothReads SLIDINGWINDOW:4:20 MINLEN:40. The filtered rRNA-free paired-end reads were aligned to the *Solanum lycopersicum* reference genome (SLM_r2.1; GenBank accession: GCF_036512215.1) using STAR^68^. FeatureCounts (version 2.0.2)^69^ was used to count reads per annotated gene for downstream differential gene expression analysis.

Differential expression analysis was performed using the R package limma^70^. For this, read counts were converted to log2 counts per million (CPM) and genes were filtered for those with at least 20 reads. The method of trimmed mean of M-values (TMM)^71^ from edgeR^72^ was then applied to calculate normalization factors and to account for library size variations. Differentially Expressed Genes (DEGs) were identified using a significance threshold (Benjamini–Hochberg adjusted *p* < 0.05) after Benjamini-Hochberg corrections. We also calculated plasticity indices (π)^73^ for each gene to assess overall gene expression bias among the treatments. All computed metrics, including logarithmic fold-changes (logFC), adjusted p-values and π for genes can be found in **Supplementary Table S6**.

For Gene Ontology (GO) enrichment analyses, we used the R package topGO ^74^ to identify biological processes significantly associated with DEGs between groups (FDR-adjusted *p* < 0.05; **Supplementary Tables S2-S4)**. Functional enrichment was performed for pairwise comparisons with sufficient numbers of DEGs, including ExBRKN versus control (360 genes), plant extract versus control (278 genes), and RKN versus ExBRKN (311 genes) (Table 1). Comparisons with only 1–4 DEGs were excluded from functional enrichment analyses.

### Figure generation and statistical analyses

All statistical analyses and graphical outputs were conducted in R (version 4.5.1)^75^ using the RStudio environment^76^. Data visualization was performed using the ggplot2 package^77^. The effects of foliar application of cavalcade on gall formation and nematode development within tomato roots were analysed separately, as these parameters showed statistically significant differences based on t-tests (*p* ≤ 0.05). In contrast, the infectivity of second-stage juvenile nematodes was analysed collectively, since no significant differences among treatments were observed (*p* > 0.05). Mean values for each experiment were then subjected to analysis of variance (ANOVA), and when significant treatment effects were observed, mean separation was performed using Duncan’s Multiple Range Test (DMRT) at *p* ≤ 0.05.

## Data availability

All data generated for this study is included in the manuscript and the Supplementary Data. The raw RNA sequencing data are available at NCBI (BioProject: PRJNA1405117; Reviewer link: https://dataview.ncbi.nlm.nih.gov/object/PRJNA1405117?reviewer=vcgi7htt4d73iq54j6g90vk622).

## Acknowledgements

The authors gratefully acknowledge Iris Riedl-Quinkertz for her expertise and access to qRT-PCR facilities, Ann Marie Waldvogel for providing the equipment used for RNA extraction, and Jan Henrik Schmidt for providing the root-knot nematode material used in the experiments. This work was financially supported by the Kasetsart University Institute for Advanced Studies (KUIAS) and the Faculty of Agriculture through the KU Brain & Man Program awarded to B.C. Additional funding was provided by an ENP grant to P.H.S. (grant no. 434028868) and by the Biodiversity Genomics Center Cologne (BioC2), funded through the University of Cologne (UoC) Forum within the Excellent Research Support Program. M.E. was supported by the Deutsche Forschungsgemeinschaft (DFG, grant no. BA 58004-1). L.I.V. was funded by the DFG Collaborative Research Centre CRC1211 (grant no. 268236062), sub-project B08 co-led by P.H.S. A.T. was supported by a doctoral scholarship from the German Academic Exchange Service (DAAD).

## Author Contributions

**N.B**. conceived the study, designed and performed all experiments, analysed the data and wrote the original manuscript. **T.H**. contributed to the experimental design for RNA extraction and qRT–PCR and revised the manuscript. **M.E**. performed the mRNA data analysis, revised the manuscript and supervised the project. **A.Sa**. supervised the study and revised the manuscript. **A.Su**. assisted with RNA extraction. **L.V**. assisted with sample preparation for sequencing. **A.T**. assisted with qRT–PCR preparation. **B.C**. supervised the project, secured funding, contributed to experimental design and revised the manuscript. **P.H.S**. supervised the project, secured funding and resources, and revised the manuscript. All authors reviewed and approved the final manuscript.

