## Supplementary material for "Cavalcade-Mediated Resistance Alters Tomato–Root-Knot Nematode Interactions and Limits Nematode Infection": Suplementary fig. and tables.docx

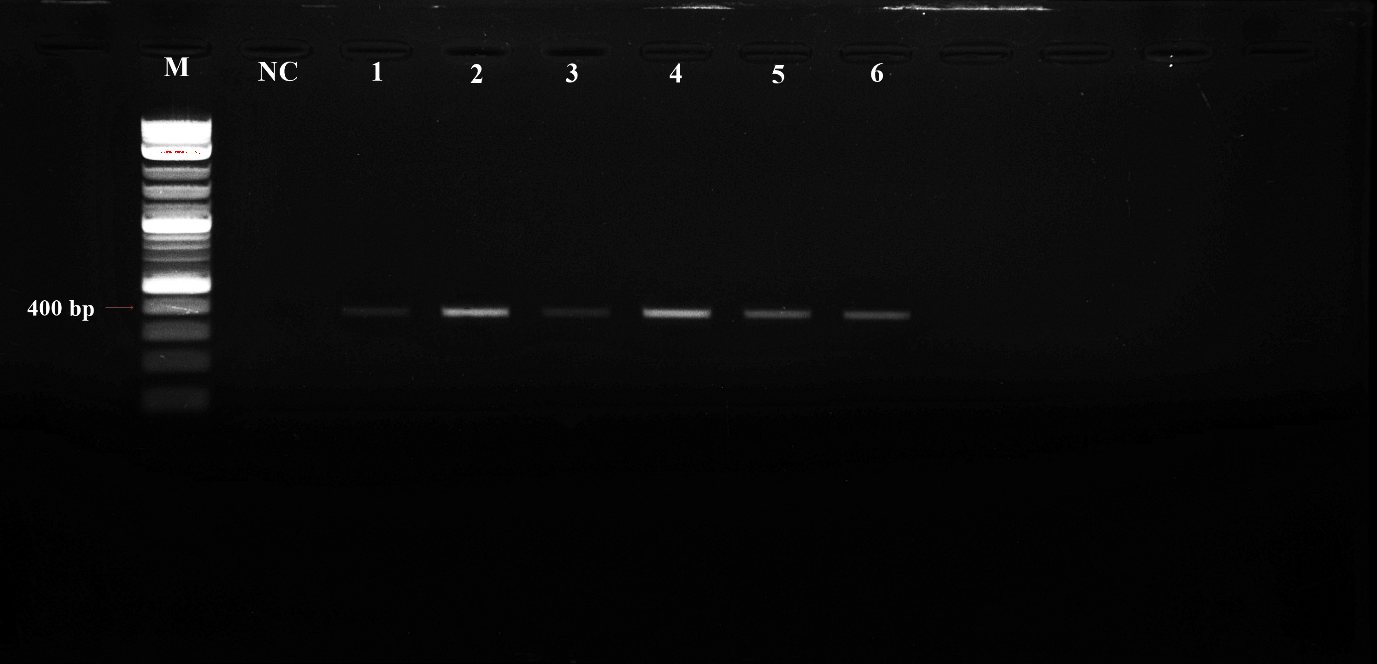

**Figure S1** PCR amplification of *Meloidogyne incognita* DNA using the SCAR marker Inc14F/Inc14R.

M, 1 kb Plus DNA ladder; NC, negative control (no template DNA); lanes 1–6, genomic DNA from *M. incognita* populations analysed in this study.

**Table S1** Effects of cavalcade leaf aqueous extract on root weight, plant height, and fresh shoot weight of tomato plants under *Meloidogyne incognita* infection in two independent trials. Data are presented as mean ± SE (n=4). Statistical differences were analysed by ANOVA.

| **Treatment** | **Plant growth parameters** | | |
| --- | --- | --- | --- |
|  | **Root weight**  **(g)** | **Plant height**  **(cm)** | **Fresh shoot weight**  **(g)** |
| **Trial 1** |  |  |  |
| DW | 0.3±0.1 | 15.2±1.5 | 4.2±1.1 |
| Extract | 0.4±0.1 | 20.9±1.5 | 6.5±1.0 |
| RKN | 0.4±0.1 | 17.1±1.9 | 4.9±1.2 |
| ExBRKN | 0.6±0.1 | 19.5±2.8 | 5.7±1.2 |
| RKNBEx | 0.6±0.1 | 21.0±1.8 | 6.6±1.2 |
| *P* value | 0.307 | 0.198 | 0.764 |
| **Trial 2** |  |  |  |
| DW | 0.6±0.2 | 21.7±2.5 | 8.1±1.9 |
| Extract | 0.6±0.2 | 25.0±0.5 | 10.1±1.2 |
| RKN | 0.7±0.2 | 24.2±1.4 | 10.0±1.8 |
| ExBRKN | 0.9±0.2 | 25.8±1.2 | 11.3±1.3 |
| RKNBEx | 0.7±0.2 | 24.7±1.8 | 10.0±1.4 |
| *P* value | 0.758 | 0.478 | 0.693 |

**Table S2** Gene Ontology (GO) enrichment analysis of differentially expressed genes between ExBRKN and distilled water treatments.

| **Upregulation** | | | | | | | |
| --- | --- | --- | --- | --- | --- | --- | --- |
| **GO.ID** | **Term** | **Annotated** | **Significant** | **Expected** | **classicFisher** | **FDR** | **ontology** |
| GO:0005385 | zinc ion transmembrane transporter activity | 18 | 4 | 0.12 | 4.80E-06 | 0.00096 | Molecular_Function |
| GO:0003700 | DNA-binding transcription factor activity | 1125 | 20 | 7.31 | 1.50E-04 | 0.015 | Molecular_Function |
| GO:0006355 | regulation of DNA-templated transcription | 1608 | 24 | 10.01 | 8.10E-06 | 0.00162 | Biological_Process |
| GO:0071577 | zinc ion transmembrane transport | 16 | 3 | 0.1 | 1.20E-04 | 0.012 | Biological_Process |
| GO:0009873 | ethylene-activated signaling pathway | 59 | 4 | 0.37 | 4.90E-04 | 0.03266667 | Biological_Process |
| **Downregulation** | | | | | | | |
| **GO.ID** | **Term** | **Annotated** | **Significant** | **Expected** | **classicFisher** | **FDR** | **ontology** |
| GO:0016762 | xyloglucan:xyloglucosyl transferase activity | 38 | 7 | 0.27 | 9.20E-09 | 0.00000184 | Molecular_Function |
| GO:0140662 | ATP-dependent protein folding chaperone | 50 | 6 | 0.36 | 1.60E-06 | 0.00016 | Molecular_Function |
| GO:0051082 | unfolded protein binding | 103 | 6 | 0.75 | 1.00E-04 | 0.006666667 | Molecular_Function |
| GO:0000234 | phosphoethanolamine N-methyltransferase | 3 | 2 | 0.02 | 1.60E-04 | 0.008 | Molecular_Function |
| GO:0003700 | DNA-binding transcription factor activity | 1125 | 17 | 8.14 | 6.80E-04 | 0.0272 | Molecular_Function |
| GO:0010411 | xyloglucan metabolic process | 48 | 9 | 0.34 | 7.60E-09 | 0.00000152 | Biological_Process |
| GO:0042546 | cell wall biogenesis | 133 | 10 | 0.94 | 1.80E-08 | 0.0000018 | Biological_Process |
| GO:0034605 | cellular response to heat | 32 | 5 | 0.23 | 2.70E-06 | 0.00018 | Biological_Process |
| GO:0006457 | protein folding | 186 | 8 | 1.31 | 9.10E-04 | 0.042571429 | Biological_Process |
| GO:0006656 | phosphatidylcholine biosynthetic process | 8 | 2 | 0.06 | 1.33E-03 | 0.042571429 | Biological_Process |
| GO:0009687 | abscisic acid metabolic process | 8 | 2 | 0.06 | 1.33E-03 | 0.042571429 | Biological_Process |
| GO:0009408 | response to heat | 66 | 8 | 0.46 | 1.49E-03 | 0.042571429 | Biological_Process |
| GO:0009969 | xyloglucan biosynthetic process | 9 | 2 | 0.06 | 1.71E-03 | 0.04275 | Biological_Process |
| GO:0048046 | apoplast | 74 | 7 | 0.51 | 7.20E-07 | 0.000144 | Cellular_Component |
| GO:0005618 | cell wall | 167 | 7 | 1.16 | 1.50E-04 | 0.015 | Cellular_Component |

**Table S3** Gene Ontology (GO) enrichment analysis of differentially expressed genes between extract and distilled water treatments.

| **Upregulation** | | | | | | | |
| --- | --- | --- | --- | --- | --- | --- | --- |
| **GO.ID** | **Term** | **Annotated** | **Significant** | **Expected** | **classicFisher** | **FDR** | **ontology** |
| GO:0009695 | jasmonic acid biosynthetic process | 6 | 3 | 0.05 | 1.50E-05 | 0.003 | Biological_Process |
| GO:0006629 | lipid metabolic process | 868 | 19 | 7.95 | 5.00E-04 | 0.04466667 | Biological_Process |
| GO:0006355 | regulation of DNA-templated transcription | 1608 | 27 | 14.73 | 6.70E-04 | 0.04466667 | Biological_Process |
| **Downregulation** | | | | | | | |
| **GO.ID** | **Term** | **Annotated** | **Significant** | **Expected** | **classicFisher** | **FDR** | **ontology** |
| GO:0034605 | cellular response to heat | 32 | 4 | 0.06 | 4.20E-07 | 8.40E-05 | Biological_Process |

**Table S4** Gene Ontology (GO) enrichment analysis of differentially expressed genes between RKN and ExBRKN treatments.

| **Upregulation** | | | | | | | |
| --- | --- | --- | --- | --- | --- | --- | --- |
| **GO.ID** | **Term** | **Annotated** | **Significant** | **Expected** | **classicFisher** | **FDR** | **ontology** |
| GO:0005388 | P-type calcium transporter activity | 16 | 4 | 0.14 | 1.10E-05 | 0.0016 | Molecular_Function |
| GO:0016762 | xyloglucan:xyloglucosyl transferase activity | 38 | 5 | 0.34 | 2.20E-05 | 0.0016 | Molecular_Function |
| GO:0005452 | solute:inorganic anion antiporter activity | 7 | 3 | 0.06 | 2.40E-05 | 0.0016 | Molecular_Function |
| GO:0005524 | ATP binding | 1916 | 34 | 17.22 | 7.70E-05 | 0.00336 | Molecular_Function |
| GO:0140662 | ATP-dependent protein folding chaperone | 50 | 5 | 0.45 | 8.40E-05 | 0.00336 | Molecular_Function |
| GO:0000234 | phosphoethanolamine N-methyltransferase | 3 | 2 | 0.03 | 2.40E-04 | 0.008 | Molecular_Function |
| GO:0005516 | calmodulin binding | 72 | 5 | 0.65 | 4.70E-04 | 0.013428571 | Molecular_Function |
| GO:0050801 | monoatomic ion homeostasis | 136 | 5 | 1.22 | 6.70E-06 | 0.00134 | Biological_Process |
| GO:0010411 | xyloglucan metabolic process | 48 | 6 | 0.43 | 2.30E-05 | 0.0023 | Biological_Process |
| GO:0042546 | cell wall biogenesis | 133 | 8 | 1.19 | 4.00E-05 | 0.002666667 | Biological_Process |
| GO:0080142 | regulation of salicylic acid biosynthetic | 12 | 3 | 0.11 | 1.40E-04 | 0.007 | Biological_Process |
| GO:0045010 | actin nucleation | 24 | 4 | 0.21 | 1.80E-04 | 0.0072 | Biological_Process |
| GO:0006820 | monoatomic anion transport | 31 | 4 | 0.28 | 5.00E-04 | 0.016285714 | Biological_Process |
| GO:0070588 | calcium ion transmembrane transport | 43 | 4 | 0.38 | 5.70E-04 | 0.016285714 | Biological_Process |
| GO:0048046 | apoplast | 74 | 6 | 0.61 | 3.00E-05 | 0.006 | Cellular_Component |
| **Downregulation** | | | | | | | |
| none | | | | | | | |

**Table S5** Functional annotation of differentially expressed genes based on eggNOG analysis.

| Gene ID | query | e value | score | Description |
| --- | --- | --- | --- | --- |
| LOC101254501 | XP_004241669.1 | 3.94e-48 | 155.0 | - |
| LOC101245195 | XP_004246568.1 | 7.84e-92 | 268.0 | - |
| LOC104646728 | XP_010319387.1 | 2.72e-42 | 139.0 | - |
| Asr4 | NP_001269248.1 | 8.34e-20 | 95.1 | ABA/WDS induced protein |
| LOC101260610 | XP_004251812.1 | 0.0 | 896.0 | Anthocyanin acyltransferase |
| LOC101259430 | XP_004236999.1 | 1.12e-265 | 728.0 | B3 DNA binding domain |
| LOC104648101 | XP_010322968.1 | 1.65e-247 | 682.0 | BAG domain |
| LOC101254583 | XP_004235966.1 | 0.0 | 1732.0 | Belongs to the ClpA ClpB family |
| CYP707A2 | NP_001362844.1 | 0.0 | 934.0 | Belongs to the cytochrome P450 family |
| DMR6-1 | NP_001233840.2 | 3.26e-254 | 696.0 | Belongs to the iron ascorbate-dependent oxidoreductase family |
| GA2ox5 | NP_001234757.1 | 2.36e-246 | 676.0 | Belongs to the iron ascorbate-dependent oxidoreductase family |
| LOC101261332 | XP_004240035.1 | 0.0 | 895.0 | Belongs to the peptidase A1 family |
| PORK1 | XP_004235511.1 | 1.31e-287 | 842.0 | Belongs to the protein kinase superfamily. Ser Thr protein kinase family |
| tXET-B1 | NP_001234213.2 | 7.92e-215 | 592.0 | Catalyzes xyloglucan endohydrolysis (XEH) and or endotransglycosylation (XET). Cleaves and religates xyloglucan polymers, an essential constituent of the primary cell wall, and thereby participates in cell wall construction of growing tissues |
| LOC101258632 | XP_004235157.2 | 1.12e-213 | 589.0 | Catalyzes xyloglucan endohydrolysis (XEH) and or endotransglycosylation (XET). Cleaves and religates xyloglucan polymers, an essential constituent of the primary cell wall, and thereby participates in cell wall construction of growing tissues |
| LOC101247110 | XP_004252273.1 | 0.0 | 1021.0 | CheR methyltransferase, SAM binding domain |
| LOC101255316 | XP_019070980.1 | 0.0 | 1164.0 | Conserved gene of |
| LOC101247750 | XP_004229409.2 | 1.61e-110 | 320.0 | Cotton fibre expressed protein |
| LOC101263598 | XP_004236333.2 | 0.0 | 994.0 | Cytochrome p450 |
| LOC101248306 | XP_004241226.1 | 0.0 | 1009.0 | cytochrome P450 |
| Prg1 | NP_001234261.1 | 3.15e-161 | 451.0 | Divergent CCT motif |
| PTI4 | NP_001334005.1 | 3.28e-166 | 464.0 | DNA-binding domain in plant proteins such as APETALA2 and EREBPs |
| ERF-D1 | XP_004237483.1 | 6.02e-217 | 598.0 | DNA-binding domain in plant proteins such as APETALA2 and EREBPs |
| LOC101253047 | XP_010312777.1 | 1.36e-121 | 350.0 | DNA-binding domain in plant proteins such as APETALA2 and EREBPs |
| LOC101259708 | XP_004251216.2 | 7.07e-274 | 752.0 | dnaJ protein |
| LOC101250661 | XP_004251272.2 | 2.31e-196 | 545.0 | Dof zinc finger protein |
| LOC101265639 | XP_004239200.1 | 0.0 | 930.0 | Domain associated at C-terminal with AAA |
| LOC101261868 | XP_004232247.2 | 0.0 | 1350.0 | Domain of unknown function (DUF569) |
| AP2d | XP_010313188.2 | 0.0 | 881.0 | Ethylene-responsive transcription factor RAP2-7-like isoform X1 |
| LOC101255164 | XP_004250959.1 | 0.0 | 1273.0 | Heat shock cognate 70 kDa |
| LOC101266155 | XP_004247199.1 | 2.06e-257 | 707.0 | heat shock factor |
| HSP90 | NP_001308492.1 | 0.0 | 1248.0 | Heat shock protein |
| LOC101264183 | XP_004234218.2 | 0.0 | 1296.0 | Heat shock protein |
| ER24 | NP_001234468.2 | 8.55e-94 | 274.0 | Helix-turn-helix XRE-family like proteins |
| HSP17.4 | NP_001234130.2 | 5.9e-103 | 298.0 | kDa class II heat shock protein-like |
| LOC101262540 | XP_004244396.1 | 2.49e-255 | 699.0 | Kinase that can phosphorylate various inositol polyphosphate such as Ins(3,4,5,6)P4 or Ins(1,3,4)P3 |
| LOC101262429 | XP_004241948.1 | 2.36e-269 | 736.0 | leucine-rich repeat receptor-like protein kinase At2g19210 |
| LOC101253979 | XP_010317814.1 | 4.99e-225 | 620.0 | Mitochondrial calcium uniporter |
| LOC101248512 | XP_010325945.1 | 3.08e-166 | 484.0 | Myb/SANT-like DNA-binding domain |
| LOC101267990 | XP_004239208.1 | 1.48e-293 | 799.0 | MYND finger |
| LOC101268703 | XP_010323783.2 | 0.0 | 2027.0 | PAN-like domain |
| LOC101264995 | XP_004233351.1 | 1.03e-199 | 551.0 | PCO_ADO |
| LOC101252337 | XP_004249003.1 | 3.16e-260 | 711.0 | Phosphate-induced protein 1 conserved region |
| LOC101266879 | XP_004249051.1 | 3.75e-79 | 234.0 | Plant non-specific lipid-transfer proteins transfer phospholipids as well as galactolipids across membranes. May play a role in wax or cutin deposition in the cell walls of expanding epidermal cells and certain secretory tissues |
| LOC101256762 | XP_004236652.1 | 1.16e-118 | 339.0 | Protein kinase C conserved region 2 (CalB) |
| LOC101267255 | XP_004244577.1 | 8.38e-152 | 427.0 | Protein of unknown function (DUF1442) |
| LOC101268751 | XP_004253017.1 | 8.88e-138 | 389.0 | resistance |
| LOC101247100 | XP_004251097.1 | 1.32e-308 | 840.0 | RING-type E3 ubiquitin transferase |
| MTSHP | NP_001233872.1 | 1.83e-142 | 402.0 | small heat shock protein |
| ERF-D5 | XP_004237144.1 | 1.94e-254 | 701.0 | Transcription factor |
| ERF84 | XP_004237817.1 | 1.21e-259 | 719.0 | Transcription factor |
| LOC101266819 | XP_004238969.1 | 2.04e-168 | 477.0 | Transcription factor |
| LOC101253074 | XP_004236453.2 | 0.0 | 1081.0 | UPF0481 protein At3g02645 |
| LOC101254813 | XP_004243310.1 | 4.44e-55 | 171.0 | Wound induced protein |
| LOC101254505 | XP_010323709.2 | 4.55e-57 | 178.0 | Wound induced protein |
